# Benchmarking and optimization of cell-free DNA deconvolution

**DOI:** 10.1101/2023.07.17.549353

**Authors:** Tom Hill, Neelam Redekar, Temesgen E. Andargie, Moon K. Jang, Sean Agbor-Enoh

**Author notes:** Address Correspondence to: Sean Agbor-Enoh, MD, PhD Lasker Clinical Research Tenure Track Investigator Laboratory of Applied Precision Omics Genomic Research Alliance for Transplantation (GRAfT) Division of Intramural Research National Heart, Lung, and Blood Institute 10 Center Dr., Rm 7D05, Bethesda, MD 20892.

## Abstract

Reference methylomes, used in deconvolution algorithms to determine cell-free DNA tissue sources, were based on driver CpGs from either microarray or sequencing platforms. Cross-validation of these algorithms is important to allow interpretation of data across studies, select optimal sequencing depth, and thus reduce costs of cf-DNA deconvolution assays. Towards this end, we assessed the performance of two reference-based deconvolution algorithms: ‘cfDNAme’, sequencing-based methylome signatures, and ‘Meth-Atlas’, a microarray-based methylome signatures using a cfDNA bisulfite sequencing. While both algorithms use NNLS model, cfDNAme uses CpG windows, while Meth-Atlas uses individual CpGs as cell or tissue signatures. We determined the optimal the number of informative CpGs signatures, and the best sequencing depths for precise deconvolution. We found that above 5-fold coverage, much lower coverage than what is frequently used, there is little difference between our two chosen algorithms, both identifying the correct tissue make-up with a high accuracy, suggesting that whole genome bisulfite sequencing for tissue of origin identification can be completed in a much more cost-effective manner than previously thought.

## 1. Introduction

Disease diagnosis often relies on less sensitive conventional clinical markers or invasive tissue biopsies ^[1]^. Recent investigation focused on more sensitive and noninvasive early diagnostic methods of tissue injury called cell-free DNA (cfDNA), which is released into the circulation following cell death from diverse tissue types ^[2, 3]^. Circulating cfDNA carries epigenetic markers, such as DNA methylation, informing its tissues-of-origin and biological pathways related to disease pathogenesis ^[4]^. DNA methylation of the cytosine at the fifth carbon position (5mC) is the most prevalent and important epigenetic mechanism. It is critical in diverse biological processes, including cell, tissue, or organ-specific gene expression, function, and identity, and it shows disease-specific alterations^[5]^. Recent studies leveraged tissue-specific DNA methylation patterns and developed deconvolution algorithms to deconvolute the tissue source of cfDNA^[5-7]^.

Owing to high base resolution, whole genomic bisulfite sequencing (WGBS) is the current standard method to capture the genome-wide DNA methylation profile. However, the analysis of WGBS presents a significant computational challenge. Currently, only few computational algorithms have been developed for cfDNA tissues-of-origin analysis generated by WGBS^[6, 8-10]^. One of these cell-type deconvolution methods is the Meth-Atlas algorithm, which employs array-based reference methylomes^[8]^, but has been used for WGBS data analysis^[11, 12]^. However, reference methylome in this approach captures less than 10% of the 28 million CpG sites of the human genome and might not be sufficient to detect low-fraction cell types accurately^[8]^. Other reference-based deconvolution methods such as cfDNAme use cell-type specific methylation windows to deduced from WGBS data^[3, 6, 9, 10]^ but might have higher noise than array data. Despite the speculation, direct comparisons of the performance of such algorithms are limited. Optimal CpG signature type and count different algorithms is required to standardize the approach and ensure reproducibility. Another technical constraint with cfDNA deconvolution analysis is the predominance of blood cells^[8]^, with nonhematopoietic tissues contributing to less than 15% of the cfDNA. Deconvoluting such low fraction tissue types may be resolved by sequencing at greater depth^[8]^ which adds to the overall sequencing cost. Thus, determining optimal sequencing depth for accurate detection of rare or low fraction cell types, is critical for deploying cfDNA technologies at reasonable sequencing costs.

This study addresses three critical questions about cfDNA methylome sequencing and deconvolution analysis: (1) What is the optimal cfDNA sequencing depth or coverage to detect low fraction cell types cost-effectively? We sequenced samples from low to high depth and examined the effects of coverage on tissue-specific cfDNA profiles. (2) What is the optimal number of informative CpG sites or windows to deconvolute different cell types and provides reproducible result? We surveyed and compared different cell specific informative CpG numbers (between 50 and 6000 sites). (3) What is the appropriate computational tool to deconvolute cell types? We compared algorithms based on whole-genome bisulfite sequencing and microarrays. We, therefore, performed whole-genome bisulfite sequencing from heart transplant patients and an artificial mixture of known cell types at different dilution fractions to evaluate sequencing depths, appropriate deconvolution algorithm to identify the tissue origin of cfDNA and informative CpG numbers.

## 2. Materials and Methods

### 2.1. Patient recruitment, samples collection, and processing

Heart transplant patients were recruited from a multi-center Genome Transplant Dynamics study (Unique identifier: NCT02423070). Peripheral blood samples were collected into Streck tubes (reference company). Healthy adult volunteer plasma samples were obtained from an NIH Clinical Center Department of Transfusion Medicine protocol (Collection and Distribution of Blood Component from Healthy Donors for In Vitro Research Use; ClinicalTrials.gov NCT00001846). Written informed consent was obtained from all participants. An overview of the experimental and computational workflow is outlined in Fig. 1A.

**Figure 1.**
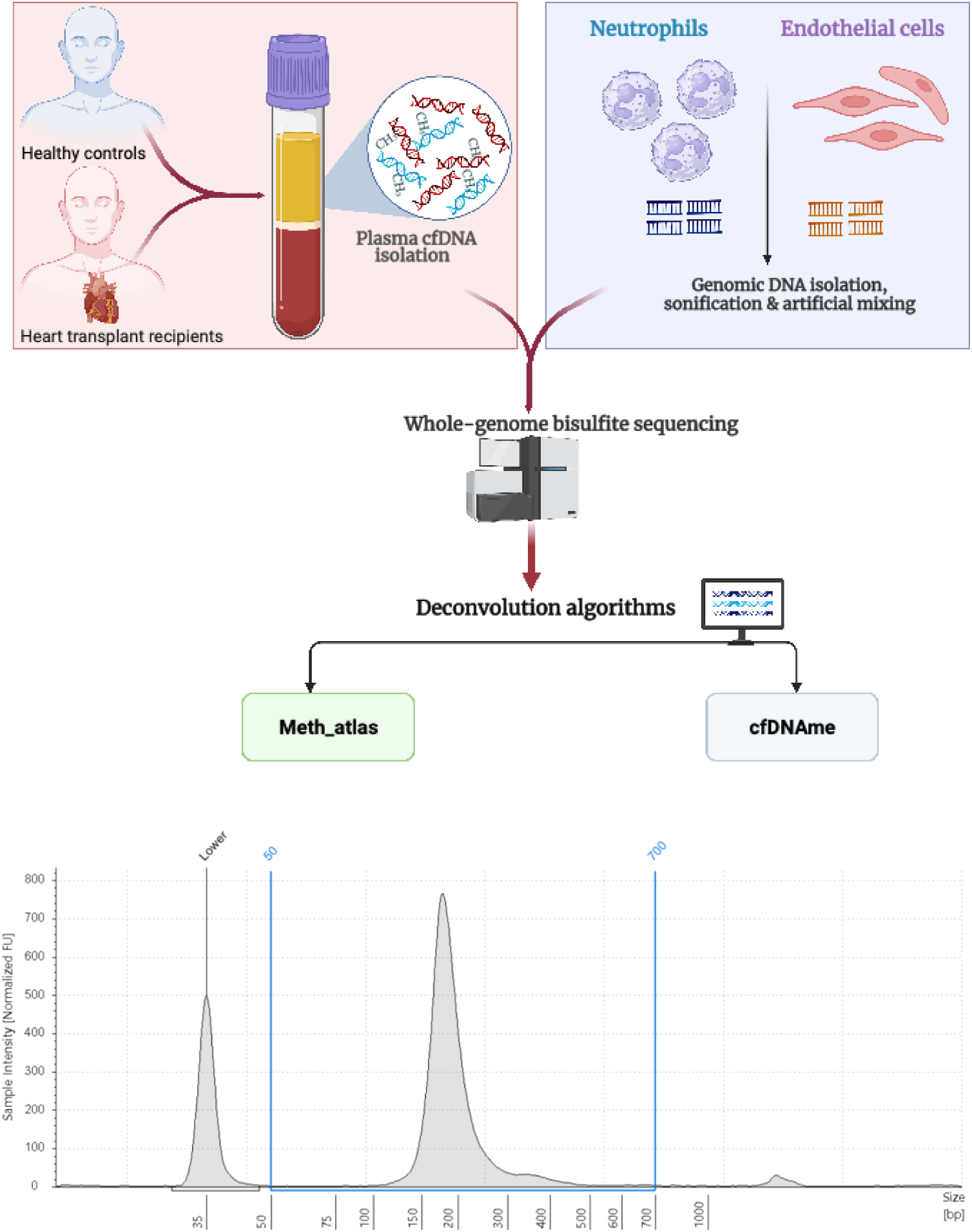
(A) Schematic demonstrating experimental workflow employed for collecting cfDNA and identifying constitutive tissue make up using methylation markers. (B) Representative plot demonstrating the cfDNA fragment size distribution.

Plasma was separated within 2hr sample collections by centrifuging at 1,600g for 10min at 4°C followed by 16,000g for 5 min at 4°C to remove residual debris. The plasma samples were spiked with 160 bp fragmented unmethylated lambda DNA (0.142 ng/mL; Promega) before cfDNA isolation to measure extraction efficiency and bisulfite conversion efficiency. Cell-free DNA was then isolated from 1 mL plasma using an automated QIAsymphony® (QS) SP Circulating DNA Kit (Qiagen, Hilden, Germany) according to the manufacturer’s guidelines, with an elution volume of 60μL. Size distribution of the cfDNA was determined with high-sensitivity Cell-free DNA ScreenTape assay according to the manufacturer’s protocol. cfDNA fragment length distributions showed a prominent peak of mononucleosome at approximately 167 bp, indicating a high-quality cfDNA profile (Fig. 1B).

Genomic DNA (gDNA) was extracted from peripheral blood samples obtained from healthy controls (n=6) and human neutrophils and human vascular endothelial cells (HUVEs) by using the QIAmp Blood Mini Kit (QIAGEN Germantown, MD) according to the manufacturer’s instructions. gDNA was sheared via sonification to ∼300 base pairs on ME220 Ultrasonicator (Covaris, Woburn, MA) and quantified by Qubit dsDNA High Sensitivity Assay Kit (Thermo Fisher). The quality of fragmented gDNA was assessed using TapeStation 2200. Each sample was then spiked with unmethylated lambda DNA (Promega) for quality control purposes or to evaluate the bisulfite conversion efficiency.

### 2.2. Total plasma cfDNA quantification

The concentration and integrity of circulating cfDNA were quantified as described earlier using short (Alu115) and long (Alu247) fragments of human Alu repetitive DNA elements (PMID: 33651717). Briefly, cfDNA samples were diluted to 10-fold in LowTE and 2 μL was used for the templated DNA in a 10 μL PCR reaction. The concentration of Alu115 reflects total cfDNA levels in the plasma (ng/mL), including both shorter and longer fragments, whereas the values of ALU247 represent longer fragments. The integrity of cfDNA is calculated as the ratio of ALU247 concentration to ALU115, which ranges between 0 and 1. The cfDNA integrity calculated from all the samples was less than 0.6, indicating proper sample collection and cfDNA isolation. The extraction efficiency of cfDNA was evaluated by dividing the concentration of lambda DNA detected after cfDNA isolation by the amount of spiked lambda DNA before cfDNA isolation. The reaction mixture consisted of 2μL cfDNA template (10-fold diluted), 5μL SYBR Green SuperMix (Bio-Rad), 1μL primer and 2μl nuclease-free water in a total of 10 µl reaction volume. The primer sequences were as follows: ALU115 (5′-CCTGAGGTCAGGAGTTCGAG-3′ [forward]) and 5′-CCCGAGTAGCTGGGATTACA-3′ [reverse]), ALU247 (5′-GTGGCTCACGCCTGTAATC-3′ [forward]) and 5′-CAGGCTGGAGTGCAGTGG-3′ [reverse]) and λ DNA (5′-GACCTCTATGCCAACACAGT-3′ [forward]) and 5′-AGTACTTGCGCTCAGGAGGA-3′ [reverse]). Each amplicon was performed in triplicate under the following conditions: initial denaturation for 5 min at 95 °C, 1cycles of denaturation at 95 °C, followed by 35 cycles of melting for 10 s at 95 °C and annealing for 1 min at 64 °C. cfDNA concentrations were expressed as ng/µL.

### 2.3. Library preparation and whole-genome bisulfite sequencing

For each sample, 5 - 50 ng of isolated cfDNA and gDNA, were used to perform bisulfite conversion using EZ DNA methylation-Gold kit (Zymo Research) as per the manufacturer’s recommendation. Libraries were generated using the Accel-NGS Methyl-Seq DNA Library Kit with Unique Dual Indexing (Swift Biosciences) for whole-genome bisulfite sequencing according to the manufacturer’s instructions. The quality of the library was assessed using a high-sensitivity D1000 ScreenTape, quantified using the Quant-iT PicoGreen dsDNA Assay kit (Life Technologies) and pooled in equimolar concentrations. The pooled library was sequenced as a 2×100 paired end on the Illumina NovaSeq 6000 platform, which resulted in low, medium, and high coverages.

### 2.4. Sequencing data processing

Following sequencing, we processed data using the methyl-seek Snakemake^[13]^ workflow (https://github.com/OpenOmics/methyl-seek). We first trimmed raw sequencing reads using TrimGalore^[14]^, and then aligned to the bisulfite converted hg38 human genome reference using Bismark version 0.23.0^[15]^. Again, we used Bismark to remove duplicate read alignments and extract the methylation status of each cytosine base in the aligned position. This step generated a CpG bed file that was further filtered to remove CpGs with low coverage or low representation across the samples within the dataset. We then converted the hg38 genomic coordinates from CpG bed file to hg19 genome build to be compatible with both deconvolution methods using the tool CrossMap^[16]^.

As the Meth-Atlas was not given as windows of the genome, but as labelled CpG markers, we used a custom python script (available at https://github.com/OpenOmics/methyl-seek) to convert the methylation status of the window overlapping each CpG to a single nucleotide-wide window to represent the CpG marker. We then used these CpG bed files with lift over coordinates as an input for deconvolution analyses using Meth-Atlas^[8]^, and cfDNAme^[10]^ methods.

### 2.5. Cell or tissue specific methylation atlas and deconvolution

For meth-atlas, we directly used the reference methylome atlas made available on the GitHub repo^[8]^. This atlas consists of ∼390,000 CpG sites which have undergone a feature selection process to subsample to 7,891 sites which are differentially expressed between each tissue type and all other tissues. For cfDNAme, we reconstructed the reference methylome atlas to include additional cell or tissue types using the Snakemake^[13]^ script provided on the GitHub repo to generate the atlas^[10]^. We downloaded the methylation profiles for additional cells and tissues (adipocytes, bladder cells, breast epithelial tissue, cortical neurons, endometrium, pancreas cells, skeletal muscle cells and vascular endothelial cells)^[9]^.

Using default script parameters provided in the cfDNAme Snakemake scripts, we used Metilene^[17]^ to compare all groups of tissues to each other in a pairwise manner and identify windows which are differentially methylated consistently between each pair of tissues (parameters: max distance between CpGs = 100, minimum CpGs to compare = 10, minimum difference in methylation = 0.2). We then extracted all genomic windows which were differentially methylated in at least one comparison, and with at least 20x coverage on average in our reference database. These high coverage cell or tissue type specific differentially methylated regions were then included in the reference methylome atlas for cfDNAme.

Before deconvolution with Meth-Atlas, we removed CpG sites with low support (less than 2-fold coverage) from each sample.

### 2.6. Effects of read coverage on cfDNA deconvolution

We generated high coverage (greater than 45x) sequencing data for our 8 ‘gold-standard’ samples (Table 1). We randomly subsampled different proportions (50%, 40%, 30%, 20%, 10%, 5%, 2.5%, 1.25%, 0.625%) of read pairs, 20 times for each of these samples. This resulted in 20 in-silico sets of read pairs at approximately 25x, 20x, 15x, 10x, 5x, 2.5x, 1.25x, 0.625x, 0.3125x coverage each for every sample. These sets of read pairs were processed to generate CpG methylation profiles and then deconvolution with Meth-Atlas and cfDNAme as described above.

**Table 1:** Summary of the coverage for high quality/high coverage ‘gold-standard’ samples used in this study.

| Sample | Total read coverage |  | Mean read coverage for signature CpG sites/regions |  | Total CpG signatures detected |  |
| --- | --- | --- | --- | --- | --- | --- |
|  | Mean | Standard Deviation | Meth-Atlas | cfDName | Meth-Atlas (of 7891) | cfDName (of 17478) |
| MKJ8231_1_S1 | 65.70 | 85.86 | 66.12 | 66.01 | 6065 | 17478 |
| MKJ8231_2_S2 | 52.66 | 75.07 | 52.87 | 52.76 | 6031 | 17478 |
| MKJ8231_3_S3 | 53.54 | 73.18 | 52.96 | 53.42 | 6030 | 17478 |
| MKJ8231_4_S4 | 53.06 | 93.08 | 53.18 | 53.17 | 6086 | 17478 |
| MKJ8231_5_S5 | 46.58 | 83.92 | 48.12 | 47.81 | 6081 | 17478 |
| MKJ8231_6_S6 | 50.87 | 89.97 | 50.91 | 50.88 | 6080 | 17478 |
| MKJ8231_7_S7 | 58.84 | 99.02 | 60.12 | 60.23 | 6084 | 17478 |
| MKJ8231_8_S8 | 59.79 | 143.11 | 61.34 | 60.92 | 6083 | 17478 |

We examined the differences in the deconvolution across different coverages considering deconvolution output from the total read coverage (with no-subsampling from the 8 ‘gold-standard’ samples, coverage greater than 45x) as the “correct” set. For the 20 replicates of each sample at each coverage level, we calculated the median cell type proportion, to give an average estimated cell content of each sample at each coverage level. We then calculated the Euclidean distance between the ‘correct’ set and each of the in-silico lower coverage set to quantify differences. Here, the Euclidean distance is the absolute difference between the gold standard estimation of each tissue proportion and the subsampled proportion, summed across all tissues, allowing us to determine changes in deconvolution across coverage:

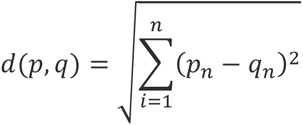

Where n represents a specific tissue, p is the estimated tissue proportion for the gold standard coverage, and q is the estimated tissue proportion in a subsampled replicate.

### 2.7. Efficacy of Meth-Atlas CpG signature markers

Efficacy of CpG signatures was estimated as the absolute difference between the mean methylation proportion of reference samples for the focal tissue (e.g., Neutrophils, noted as n), and the mean methylation proportion of samples from all other tissues (e.g., the mean methylation proportion of all non-Neutrophil samples in this case, noted as m).

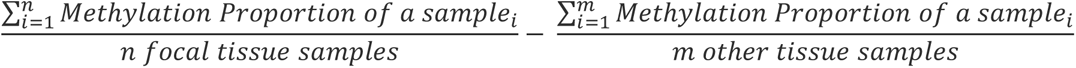

This gives an effect size, or specificity value, between 0 and 1, where 0 represents a CpG site that is no different from the background for this focal tissue (e.g. 100% methylated in all Neutrophil samples & 100% methylated in all other samples), while 1 represents a CpG site that is completely differently methylated in the focal tissue compared to the background (e.g. 0% methylated in all Neutrophil samples & 100% methylated in all other samples).

We subsampled the Meth-Atlas to the top 7000, 6000, 5000, 4000, 3000, 2000, 1000, 500, 250, 100, 90, 80, 70, 60 & 50 CpG sites for effect size (so retaining only the highest X CpG markers by effect size/specificity, before accounting for missing CpG markers which have no sequencing coverage). We then used these refined atlases to rerun deconvolution for our greater than 50x coverage ‘gold-standard’ samples. Following this, we assessed what tissues were represented in each subset of the atlas and found the Euclidean distance between the deconvolution results of the total atlas, and for these refined subset atlases.

For our healthy control and transplant sample sets, we also calculated the prevalence of CpG markers in each group (what proportion of the group each CpG site is present in) and compared this to the effect size. We also assessed how many of the CpGs for each effect size cutoff are found in each group.

### 2.8. Statistics

We performed Wilcoxon Rank Sum tests, fit generalized linear models, and calculated Euclidean distance and variance using the R statistical software^[18]^. Spearman’s Rank Correlation was used for exploring correlations. We used GGplot2 R package to generate figures in the manuscript^[19]^.

## 3. Results

### 3.1. Tissue or cell type specific reference methylomes in Meth-Atlas and cfDNAme

We first sought to assess the accuracy and reliability of a deconvolution algorithm, Meth-Atlas – that uses microarray-based methylation library, to determine tissue of origin in cell-free DNA (cfDNA) samples^[8]^, and compare with cfDNAme^[10]^deconvolution algorithm which relies on sequencing-based methylation library. Meth-Atlas and cfDNAme are reference methylome-guided classical deconvolution algorithms that use non-negative least square linear regression (NNLS) model to identify cell or tissue of origin for a cfDNA. Meth-Atlas reference methylome is composed of 7,891 differentially methylated CpG sites covering ∼4000 genomics regions; whereas cfDNAme reference methylome is composed of 17,478 cell or tissue type specific differentially methylated regions (1.609 CpGs per region on average), rather than individual CpG sites (at 5-fold coverage cutoff for the hg38 reference genome). The CpG signatures (sites/regions) within each of two reference methylomes were originally included for their unique hypermethylation or hypomethylation status in specific cells or tissue types. Differences in number and type of CpG signatures between two methods are mainly resulting from CpG sites targeted by their respective methylation platforms – Infinium EPIC array (Meth_Atlas) targeting over 935,000 CpGs vs. whole-genome bisulfite sequencing (cfDNAme) targeting millions of CpGs across the genome. Another source of variation in reference methylomes is use of different tissue or cell types for generating the reference methylomes (Table 2). While each tissue makes up roughly 5% of the markers in Meth-Atlas, regions unique to tissues make up from 0.1% to 20% of the database used for cfDNAme.

**Table 2:** Reference methylome signatures used by cfDNAme and Meth-Atlas deconvolution algorithms.

|  | cfDName |  |  | Meth_Atlas |
| --- | --- | --- | --- | --- |
| Tissue Type<sup>ϕ</sup> | Total DMRs | Avg. CpGs in signature DMRs | Size of signature DMRs (bp) | Total CpG sites |
| Adipocytes | 643 | 1.641 | 529678 | 284 |
| B-cells | 413 | 1.537 | 291290 | 329 |
| Bladder | NA | NA | NA | 292 |
| Breast | 1922 | 1.842 | 1973081 | 298 |
| CD4T-cells + CD8T-cells | 19 | 1.468 | 12086 | 385 |
| Colon epithelial cells & upper gastrointestinal tract | 1569 | 1.764 | 1440622 | 523 |
| Cortical neurons & dendritic cells | 3559 | 1.953 | 2648350 | 318 |
| Eosinophils | 166 | 1.611 | 131796 | NA |
| Erythroblasts | 24 | 1.374 | 24734 | NA |
| Head, Neck and Larynx | NA | NA | NA | 267 |
| Heart cells | 420 | 1.646 | 351523 | 282 |
| Hepatocytes | 1546 | 1.744 | 1223905 | 315 |
| Kidney cells | 566 | 1.696 | 460106 | 299 |
| Lung cells | 36 | 1.589 | 45511 | 290 |
| Monocytes & Macrophages | 51 | 1.277 | 22039 | 272 |
| Neutrophils | 142 | 1.518 | 107510 | 289 |
| NK-cells | 352 | 1.619 | 295919 | 293 |
| Pancreatic cells | 283 | 1.646 | 1302989 | 741 |
| Progenitor cells | 18 | 1 | 729 | 248 |
| Prostate | NA | NA | NA | 278 |
| Skin cells | 351 | 1.613 | 429351 | NA |
| Spleen | 16 | 1.432 | 27696 | NA |
| Thyroid | NA | NA | NA | 279 |
| Uterus and Endometrium | 3039 | 1.757 | 2035067 | 300 |
| Vascular endothelial cells | 434 | 1.685 | 342012 | 275 |

‡ Multiple tissues/cell types may share CpGs or overlapping regions. Because of this, total number of markers or regions in each method are not equal to the total numbers indicated in text.

We found only 84 overlapping windows between the reference methylomes for cfDNAme and Meth-Atlas (Supplementary Table 1) and most overlapping windows were found in tissue/cell types that are less predominant in healthy controls, suggesting that two methods are using quite different reference methylomes information that may contribute to differences in deconvolution results.

**Supplementary Table 1:**
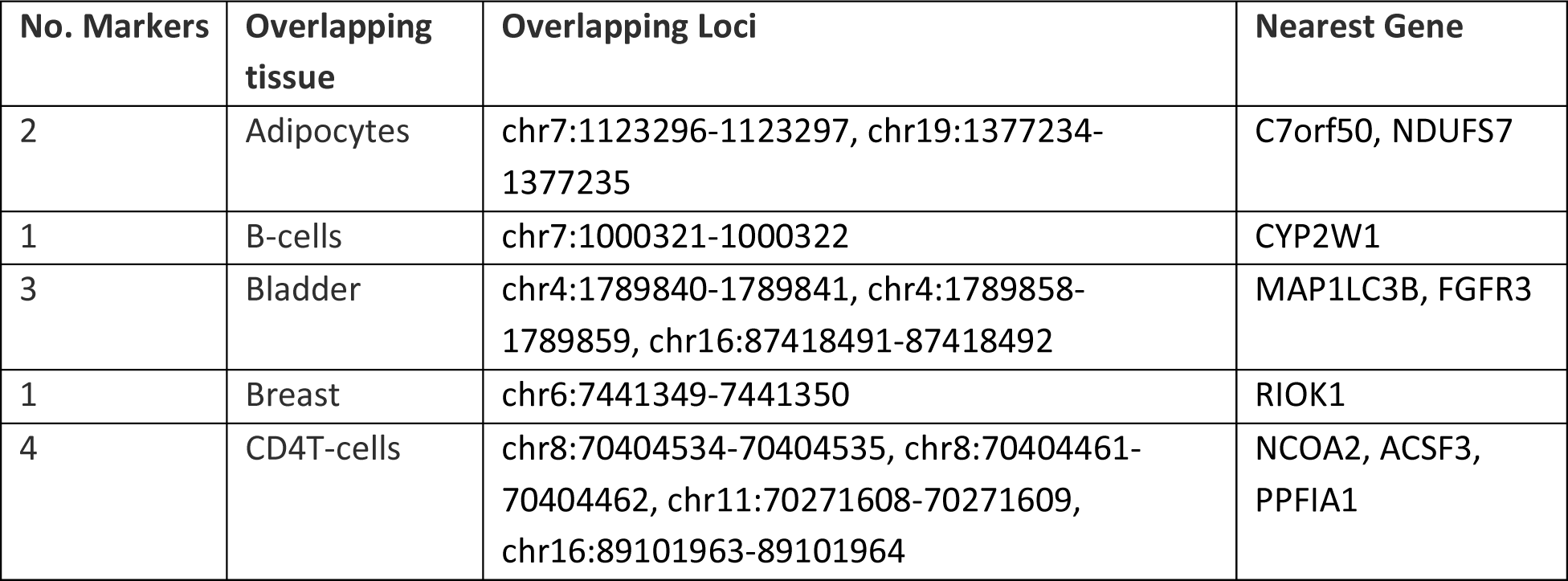
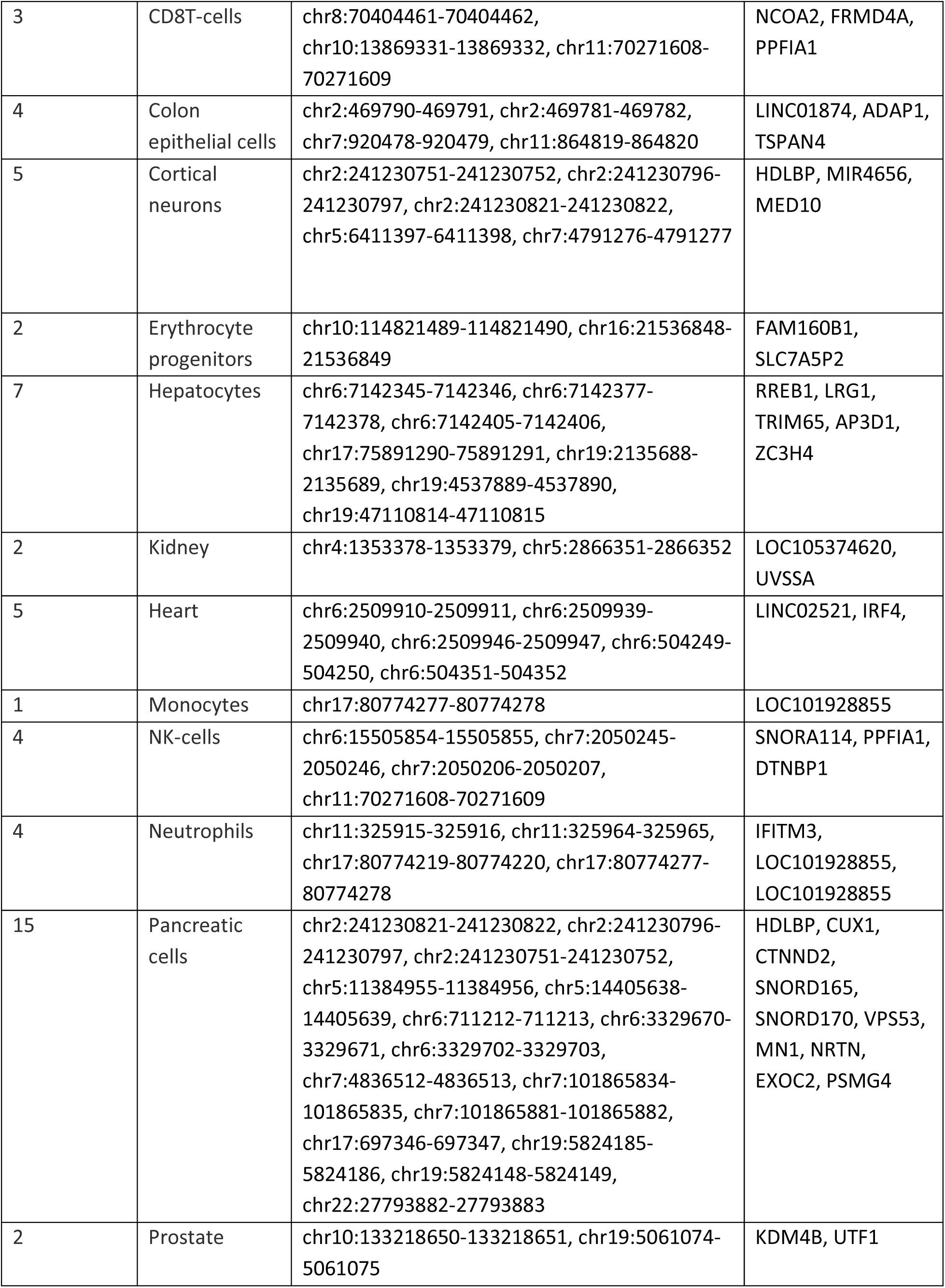
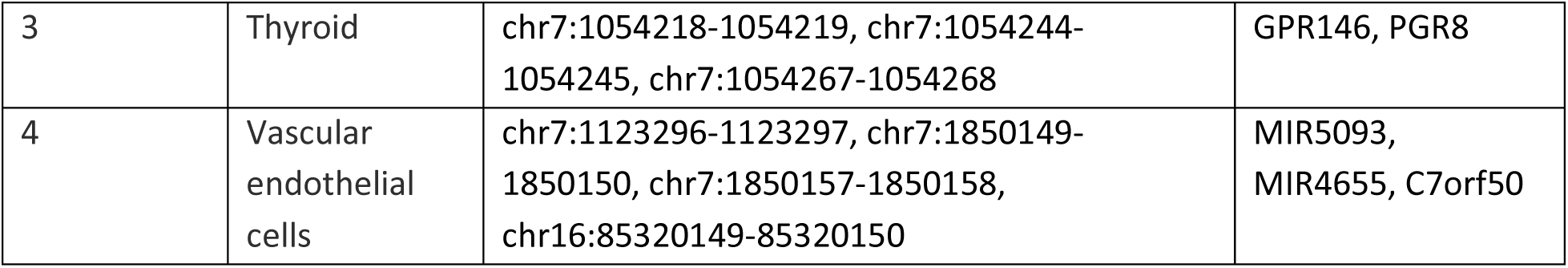
Overlapping loci between cfDNAme and Meth-Atlas databases.

Given these differences in the reference methylomes, one could argue, use of Meth-Atlas method for deconvolution of whole genome bisulfite sequencing (WGBS) data is inappropriate. Therefore, we decided to assess the accuracy and precision of Meth-Atlas with WGBS data and determine if microarray-based reference methylome is accurate enough for tissue or cell type deconvolution. We also compared Meth-Atlas and cfDNAme results with our standard samples (with known cellular composition) sequenced at varying different depths.

### 3.2. Meth-Atlas can predict accurate proportions of tissue-of-origin in WGBS data

First, we tested if Meth-Atlas could detect all cell fractions from an artificial DNA mixture of human vascular endothelial cells (HUVECs) and neutrophils. DNA isolated from these primary cells, neutrophils and HUVECs, were mixed in dilutions of 0.1 and 0.0001 dilutions with one another. The DNA mixture was subject to WGBS to an average depth of 160 million pair-end reads. The deconvolution algorithm detected DNA from both cell types down to fractions of 0.001 (1 part in 1,000 of one another DNA) with good performance. Linear regression analyses revealed the expected and observed fractions of the HUVECs (Figure 1A, R^2^ = 0.97, p < 0.001) and neutrophils (R^2^ = 0.99, p < 0.001) were strongly correlated (Figure 2A).

**Figure 2.**
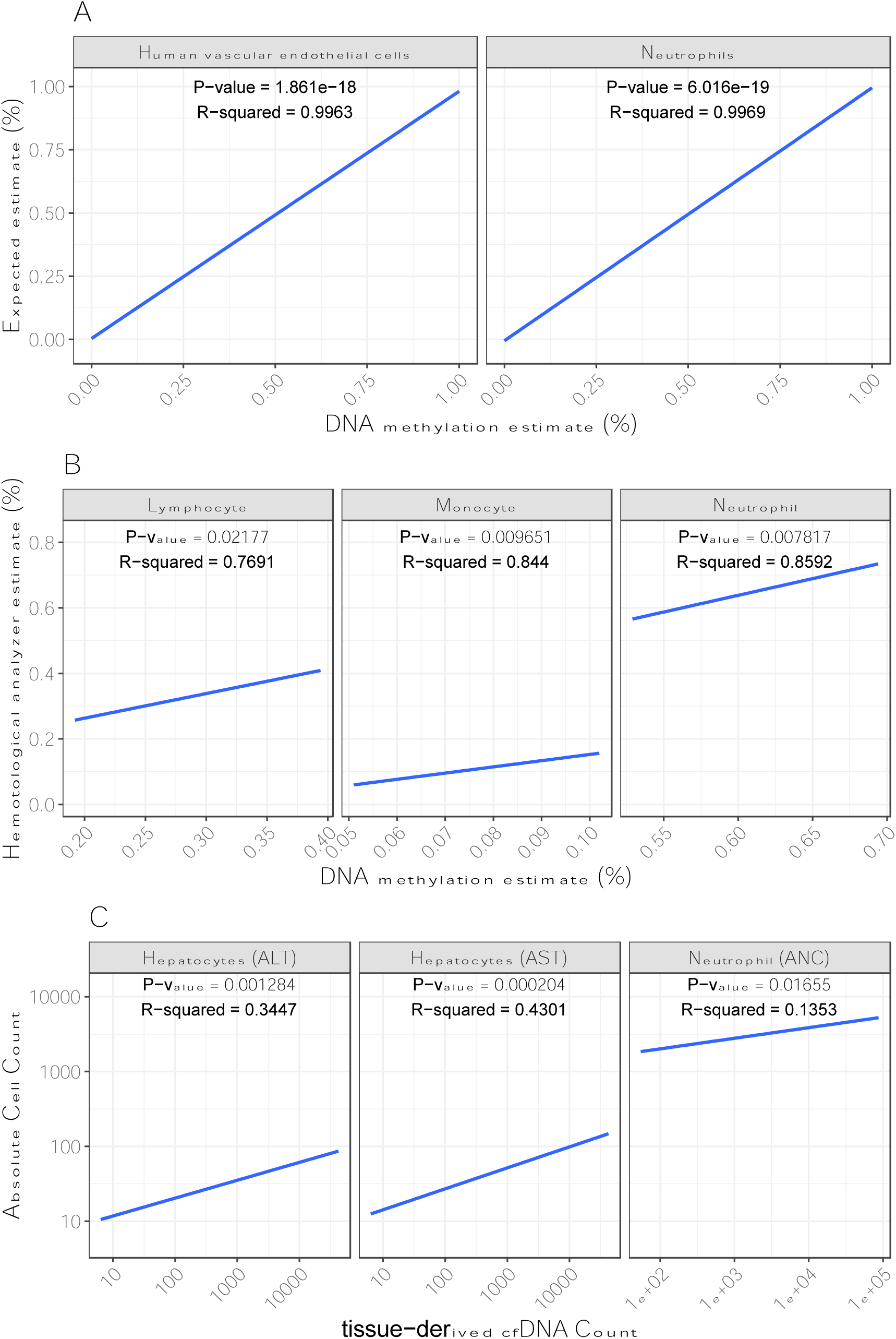
Validation of meth-atlas deconvolution algorithm. (A) Genomic DNA (gDNA) mixtures from neutrophil and human vascular endothelial cells. (B) Correlation between DNA methylation and automated hematological analyzer for leukocyte differential (granulocytes, lymphocytes, and monocytes) estimate. (C) Correlation between cfDNA and conventional clinical markers.

Next, we assessed the performance of this method to determine the fraction of leukocyte subsets in peripheral blood mononuclear cells. For this, DNA was isolated from whole blood and subjected to WGBS and deconvoluted to determine the cell-fractions. Our results indicated that the leukocyte estimates from Meth-Atlas deconvolution were highly correlated with the conventional hematological analyzer differential count, demonstrating the utility of this assay for estimating the fraction cells from a mixture of cells (Figure 2B).

Finally, we sought to investigate if tissue cfDNA measure is reflective of tissue injury and correlate with known biochemical markers of end-organ injury (Figure 2C). In heart transplant patients, we observed significant correlations between neutrophil-derived cfDNA vs. peripheral blood neutrophil counts, and liver-derived cfDNA vs. alanine transaminase (ALT) and aspartate aminotransferase (AST) liver enzymes. Collectively, these results highlight that Meth-Atlas deconvolution algorithm produces reliable and accurate results for WGBS approach. We also repeated this analysis using cfDNAme and again find the results are highly accurate (Supplementary Figure 1).

### 3.3. Meth-Atlas deconvolution accuracy remains unaffected when more than 90% of CpGs are removed

More than 77% (6,086 of total 7,891) of CpG signatures from Meth-Atlas reference methylome are used to deconvolute one type of tissue or cell (Table 2), so we assessed the contribution of different CpGs towards deconvolution accuracy. Methylation status of CpG sites varied strongly in their specificity to certain tissues, and CpG sites which were highly specific to a tissue contributed more towards the deconvolution than CpGs with less specific methylation states. We determined the effectiveness of CpG markers to deconvolute was based its specificity to a specific tissue or cell type and detection frequency.

To test this hypothesis with ‘gold-standard’ samples, we first ranked CpG markers by specificity, excluded CpG markers with low specificity, and then performing deconvolutions based on systematically reduced CpG markers from 6000 to 50 sites. For each decreasing subset of CpG markers, we used Meth-Atlas to deconvolute our gold standard samples and calculated the Euclidean distance between the results of each subset of markers and the results from all CpG markers (including ones with low specificity).

We found weak negative relationship between Euclidean distance and number of CpG markers for subsets with CpGs ranging between 6000 and 250 sites (Figure 2, GLM t-value = -2.959, p-value = 0.0436); while subsets with less than 250 CpG markers showed significant increase in the Euclidean distance (Wilcoxon Rank Sum Test, W = 2946, p-value = 1.259e-09), supporting a stronger negative relationship (Figure 2, GLM t-value = -4.272 p-value 3.95e-05). This drastic shift in Euclidean distance for these subsets used is because of the elimination of CpG markers that were specific to tissue or cell types that dominated our samples. Nonetheless, these analyses suggested deconvolutions accuracy is driven by a few highly specific CpG markers, and reduced number (>250) of specific CpG markers could still produce highly similar deconvolution results.

**Figure 2:**
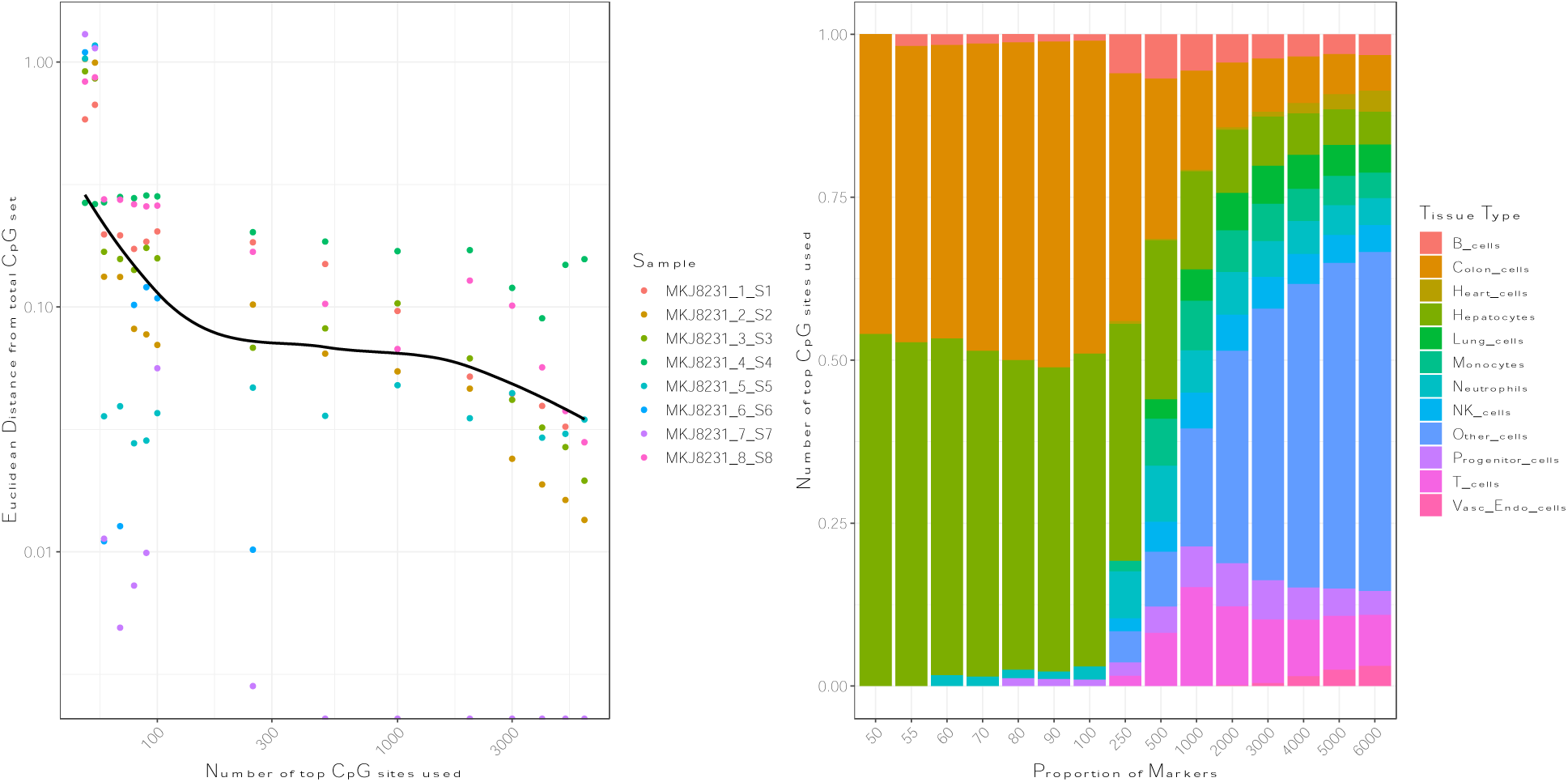
A: Effect of reducing CpG markers within reference methylome on deconvolution with Meth_Atlas algorithm. This relationship is quantified as ‘Euclidean’ distance of deconvolution results using subsets of CpG markers (as reference methylome) from the deconvolution results using entire reference methylome database. **(A)** The CpG markers were ranked by their effect size and multiple sized subsets of top-ranked CpG markers were used for deconvolution with Meth_Atlas. The Euclidean distance increases marginally as CpG markers are removed, up to the top 250 CpG markers. In subsets smaller than 250 CpGs, markers for most tissue or cell-types are missing resulting in an exponential increase in Euclidean distance. The line shows a smoothed Loess regression of the relationship between Euclidean distance & the number of top CpG markers. **(B)** Bar plot showing different tissues represented in the subsets of top CpG markers by effect size. Key tissues such as neutrophils are missing in a subset below 250 CpGs. Note that due to their small proportion, we have merged Uterus cells, Gastrointestinal cells, Bladder cells, Breast cells, Pancreas cells, Neuron cells, Adipose cells, and Skin cells as ‘Other cells’.

We also found strong linear correlation between specificity and prevalence in two sets of cfDNA sample groups (healthy and AMR) (**Supplementary Figure 2**, GLM specificity ∼ prevalence + group + group x prevalence; t-value = 79.134, p-value<2e-16, group t-value = -1.224, p-value = 0.2208).

We also assessed the number of CpGs which were shared between two groups (healthy control samples and transplant rejection samples) and found a strong overlap of highly specific CpG markers at every subset, with fewer than 10% of CpGs being found unique to each group (Supplementary Figure 3). This suggests that most CpGs aren’t group or condition specific but are generally consistently found in tissues across groups and treatments and are generally the same in tissues of different states.

### 3.4. cfDNAme and Meth-Atlas deconvolution results highly comparable despite different reference methylomes

We chose to compare Meth-Atlas to cfDNAme (an NNLS based deconvolution algorithm that utilizes reference methylome originating from WGBS data) by testing them on artificially formulated human vascular endothelial cells (HUVECs) and neutrophils samples. We found a strong positive correlation between the proportions of tissues found in each sample, across all samples (GLM: Meth-Atlas Proportion ∼ cfDNAme Proportion + sample + tissue, t-value = 34.13, p-value = 6.09e-45), with very minor differences in proportions depending on tissue composition of our sample. The most notable difference between the two databases was that neutrophils were found to be a lower proportion of the sample in cfDNAme than Meth-Atlas, while progenitor cells have a higher proportion (Figure 3, Supplementary Figure 4). As progenitor cells have very few unique CpG markers and a lot of these overlap with neutrophils & macrophages (Supplementary Figure 5, 66 of 248 Neutrophil DMR regions overlap), it is possible that these are being assigned as progenitor cells in one database, but neutrophils in the other. Further examination confirms this misassignment in cfDNAme could likely be due to a lack of high confidence CpG markers for progenitor cells (Table 2, 18 DMR regions), while this was not the case for Meth-Atlas (248 DMR regions). Similarly, we find some samples have cells misassigned as macrophages instead of Neutrophils, due to 43% of markers overlapping between the two cell types (Supplementary Figures 4 & 5).

**Figure 3:**
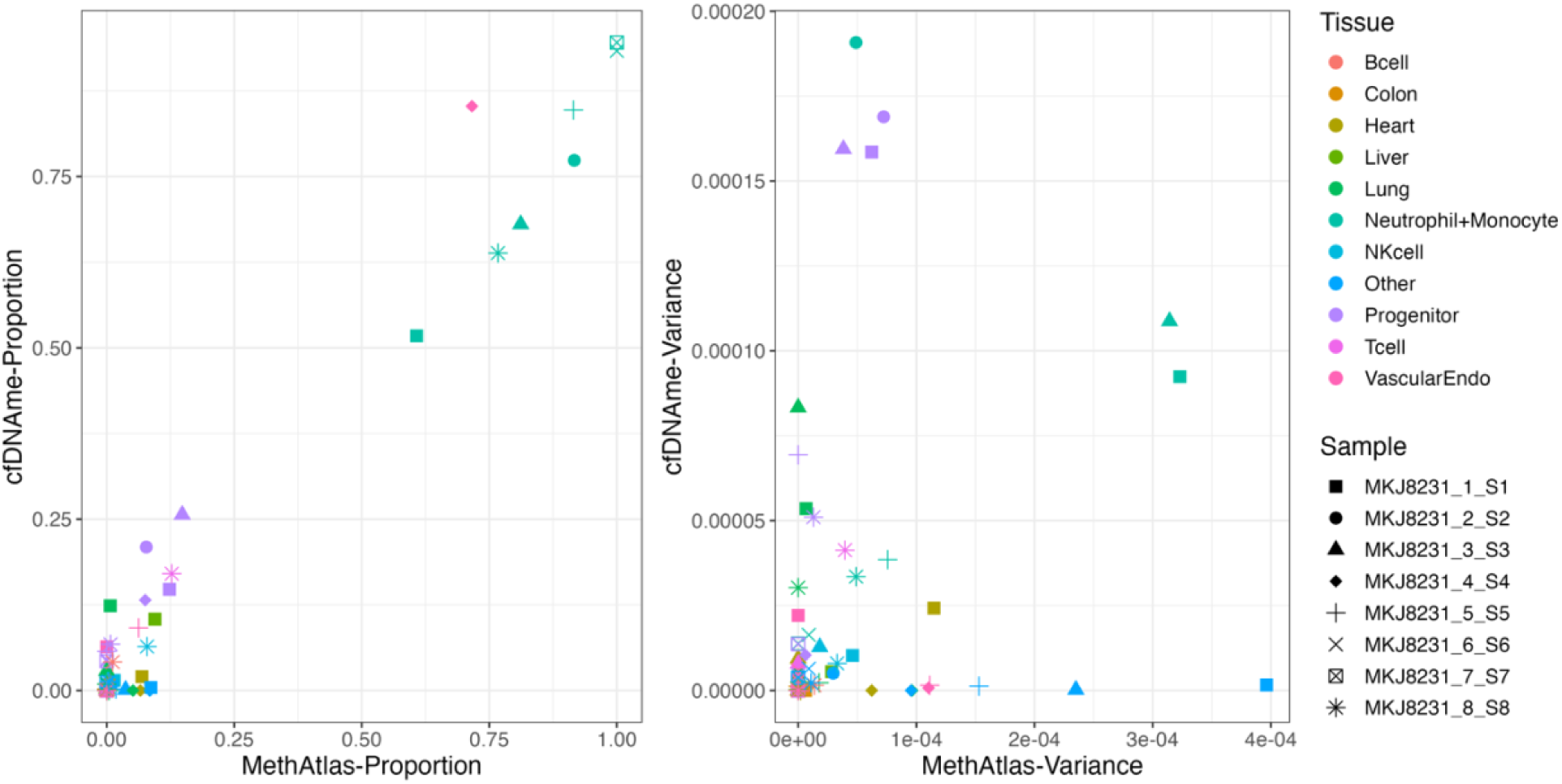
**A.** Proportion of each tissue identified for each sample, comparing the proportions estimated from cfDNAme to Meth-Atlas. **B.** The variance of estimated tissue proportions across sequencing lanes for each sample, comparing the variance of cfDNAme estimates to Meth-Atlas estimates.

### 3.5. Both cfDNAme and Meth-Atlas can accurately detect tissue types in lower sequencing depth

We further evaluated the performance of the two methods across technical replicates originating from sequencing our ‘gold standard’ sample libraries across four separate lanes of sequencing (Supplementary Figure 6). We found minor differences across lanes and between methods. For cfDNAme, the variance in proportion of ‘erythrocyte progenitor cells’ across lanes was much higher than in Meth-Atlas (Figure 3). For Meth-Atlas, the variance of rare tissues (frequency less than 1% or absent in all samples) was higher than cfDNAme, likely as rare sites were misassigned due to low coverage and so few CpG markers present which accurately represented them (Supplementary Figure 2 – specificity vs prevalence). In both cases, we found the variance for Neutrophils and Monocytes across lanes for several samples was higher than all other tissues, due to their high proportions and because both methods misassign Monocytes as Neutrophils and vice-versa across lanes, because of shared CpG markers (Supplementary Figure 5). Additionally, though sequencing depth differences between lanes was minimal, it was still present in every sample and may be contributing to the differences seen between lanes.

We next assessed the performances of Meth-Atlas and cfDNAme to deconvolute our ‘gold standard’ samples using whole-genome bisulfite sequencing approach at varying coverage levels. To do this we randomly down sampled bisulfite sequencing reads from our gold standard samples to different fractions of coverage (2.5-50%), repeated downsampling 20 times per coverage level to generate 20 sets of replicates at each coverage level. We then used cfDNAme and Meth-Atlas deconvolution methods to assign tissue types. In both cases, the median deconvolution proportions for each cell/tissue type across all replicates appeared to be similar for all subsamples with greater than 5-fold (10%) coverage (Supplementary Figure 7), though below this coverage level discrepancies start to appear between methods.

We then quantified the differences in deconvolution results for each subsampled dataset from unfractioned sample (100% coverage) across replicates, measured as Euclidean distance. We found that the Euclidean distance drops to 0 in both methods when coverage was >10% (5-fold, Figure 4) in all samples, although distance between the subsamples & the unfractioned sample was much higher in Meth-Atlas than cfDNAme as coverage gets <5-fold, likely because the Meth-Atlas deconvolution method disregards sites with lower coverage.

**Figure 4:**
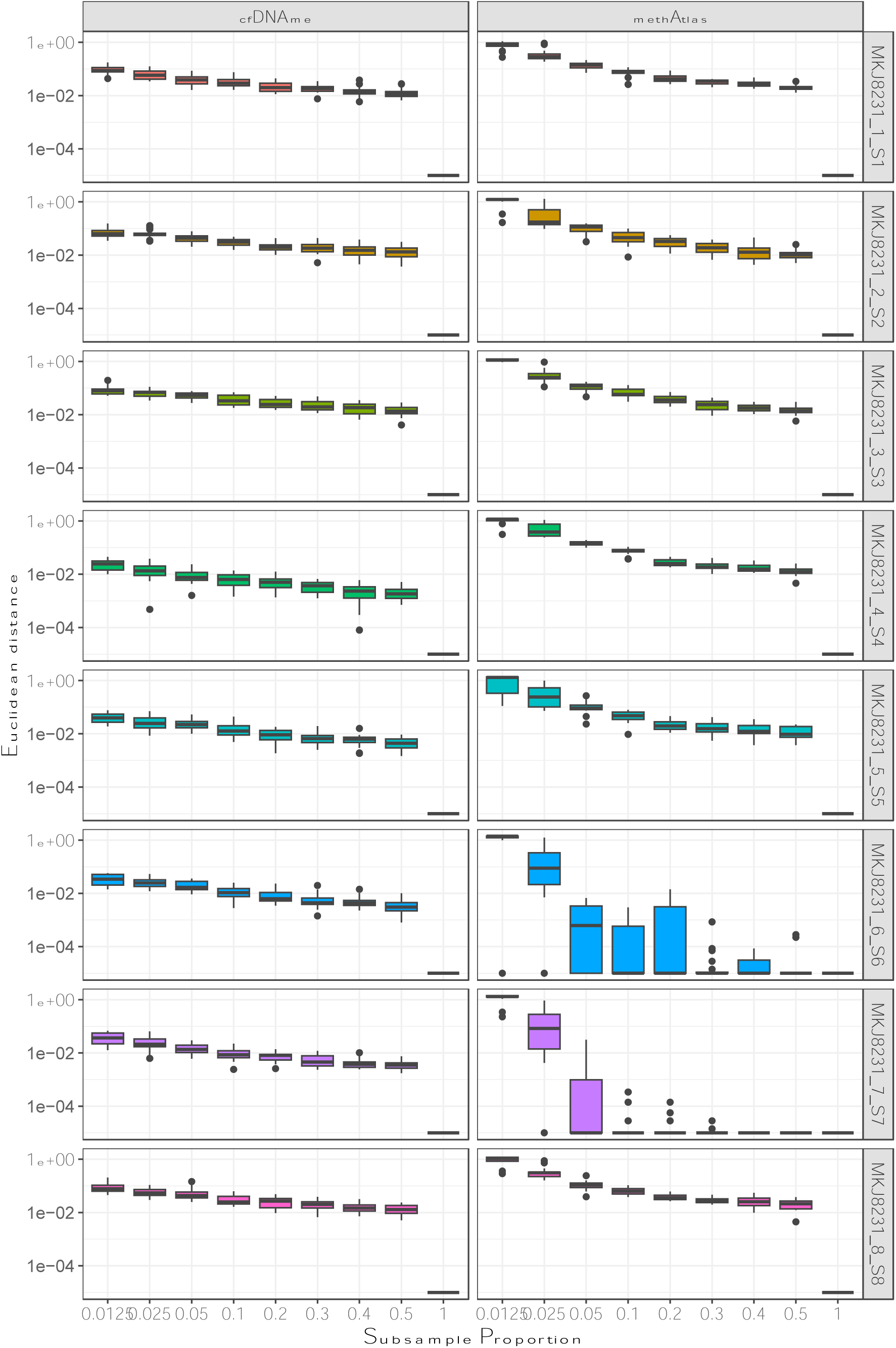
Euclidean distance of cell type proportions between subsampled replicates and the total > 45x coverage dataset, grouped by subsample proportion. Samples separated by algorithm & sample. Note that a Euclidean distance of 0 cannot be shown on a Logarithmic scale, so is shown as 1e-05.

We also assessed the variance between replicates at each subsampled coverage and find, consistent with these previous results, that variance decreases as coverage increases, but is close to 0 in all samples when 10-20% of the reads or more are used (> 5-fold coverage, Figure 5).

**Figure 5:**
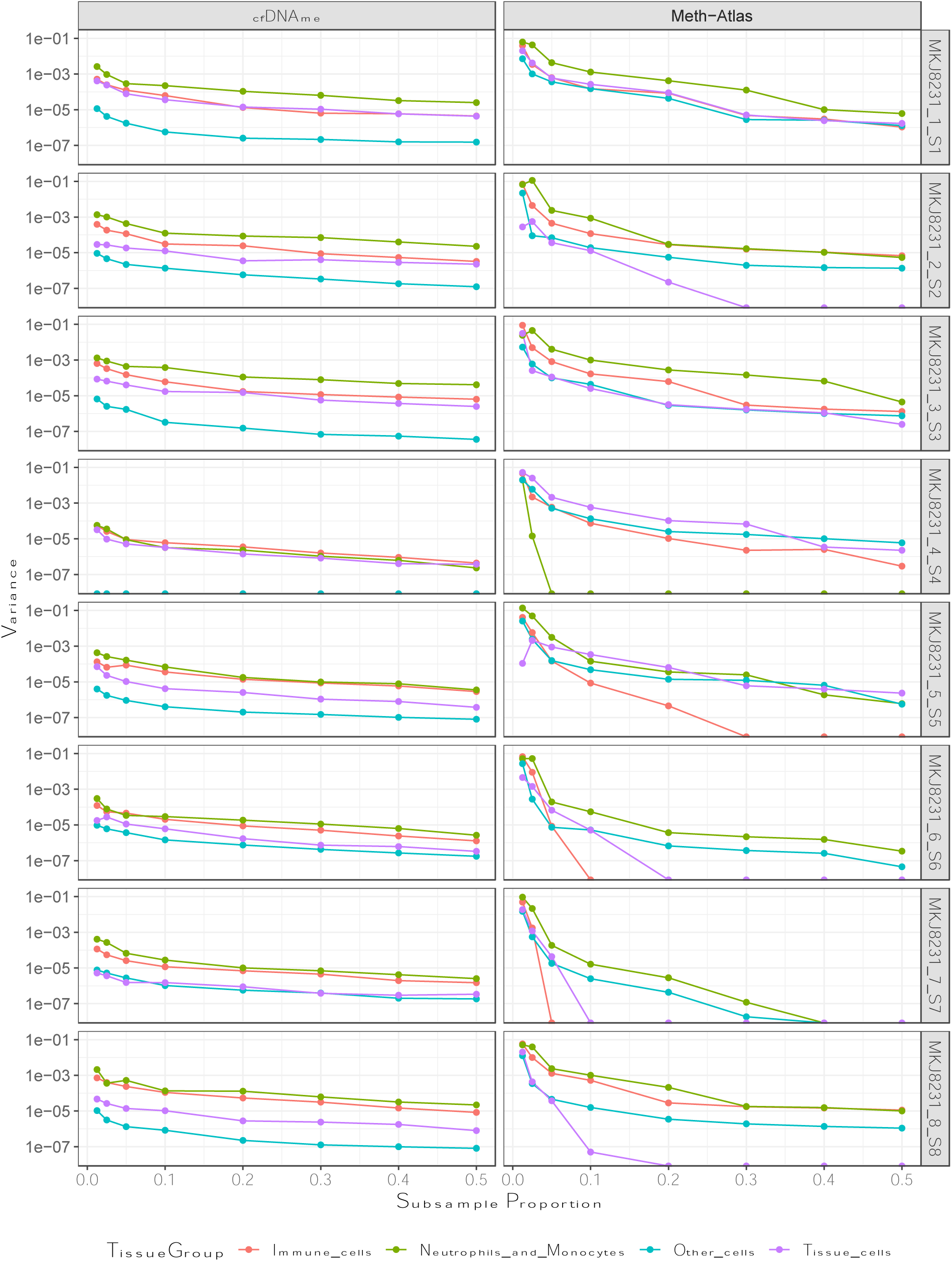
Variance across replicates for each subsampled proportion of reads, plotted separately for each sample and deconvolution algorithm. Tissues were grouped by similarity of type and proportion seen in the samples: Immune cells – T cells, B cells, NK cells, progenitor cells. Major tissue cells – Heart cells, liver cells, lung cells, Endothelial cells. Other tissue cells – Pancreas cells, Kidney cells, Bladder cells, Endometrium cells, Breast tissue cells, Skeletal muscle cells, colon cells, neuron cells, adipocytes. Note that a variance measure of 0 cannot be shown on a Logarithmic scale, so is shown as 1e-08.

A minimum coverage may be required to have an adequate number of the 7,891 CpGs to properly determine the tissue of origins of each sample, which may explain why variance levels out as coverage increases. We find a large jump in the number of CpGs found in our subsampled datasets between 5% and 20% (Supplementary Figure 8, the equivalent of 2.5-fold coverage and 10-fold coverage), above 10% (5-fold coverage) we see diminishing returns on the improved success of each deconvolution algorithm, likely because at coverage levels higher than this, most samples will have an adequate number of CpG sites (more than 500 of the most important CpG sites) to properly discern the cell types, with more CpG sites not providing any more information (Figure 2, Supplementary Figure 8).

## 4. Discussion

Most current methods for cell deconvolution of whole genome bisulfite sequencing data (WGBS) uses a non-negative least square algorithm to bin bisulfite-sequenced reads by the similarity of their methylation profile to previously determined databases of methylation calls for different tissues. This methylome database can be generated from multiple different types of data, such as using array-based data, or using WGBS data, though there has been no comprehensive examination of the difference in results seen between each database type. Here we have assessed the ability of two deconvolution databases to accurately deconvolute methylation calls from WGBS data at varying coverage levels: Meth-Atlas, which uses an array-based database, and cfDNAme, which uses a user generated database from methylation differences between a set of WGBS sequenced tissue samples. This study highlights similarities and differences between Meth-Atlas and cfDNAme deconvolution results using a range of controlled samples with known proportion of DNA mixtures from one or more cells; healthy and transplant patient samples with unknown proportion of cfDNA from different cell or tissue types. We further examined deconvolution performances at varying sequencing depth or read coverages to determine minimal sequencing data required for reliable deconvolution and explored the possible causes of differences in accuracy seen at differing coverage levels.

We have found that both methods are able to accurately determine the tissue make up of samples at higher coverages, with decreasing accuracy as coverage decreases. There is little difference between Meth-Atlas and cfDNAme algorithms at coverages greater than 5-fold average genomic coverage in samples (Figures 3 & 5). As coverage decreases, fewer CpG sites are covered and can be used to determine tissue of origin, however, we found that the Meth-Atlas was still highly accurate when using only 10% of the CpG markers, and the CpG markers which have a larger effect size are more likely to be found even at lower coverages (Figure 4 & 5). Together, these results suggest that a both cfDNAme and Meth-Atlas deconvolution methods can be used to determine tissue-of-origin accurately and reliably in cfDNA samples sequenced at low coverage in a cost-effective manner.

We suspect that there are several reasons for the minimum threshold being required for Meth-Atlas to detect the correct proportions of tissues in each cfDNA sample. Firstly, Meth-Atlas has a user given cut-off in the WGBS data (here set as 5-fold coverage) where CpG sites are not considered below this threshold. As most sites used for deconvolution are going to be either fully methylated or unmethylated, realistically we could potentially use all sites with at least 1 bisulfite converted read covering them, potentially improving the resolution of our tissue of origin estimations. However, current estimates for error rates in bisulfite sequencing are thought to be between 0.09% and 6.1%^[20]^, much higher than current error rates seen in Illumina whole-genome sequencing^[21]^. Further, higher coverages are generally used when attempting to call nucleotide variants from Illumina sequencing, which in some ways is analogous to identifying bisulfite converted sites when aligning WGBS^[15]^.

Secondly, we found that above 5-fold coverage (10% coverage in our subsampled datasets), there is a jump in the number of CpG sites covered, with coverage higher than this showing diminishing returns of the number of CpGs covered and the improvement in accuracy (Figure 2, Supplementary Figure 8). Altogether this also suggests that sequencing coverage higher than 10x is not entirely necessary for most methylation sequencing experiments involving deconvolution, much lower than previously thought.

The differences between cfDNAme are likely due to differences in our CpG reference atlas’s, rather than differences in the deconvolution algorithm. Though we have attempted to create a cfDNAme atlas which matches the Meth-Atlas tissue database, as shown in Table 2, there are numerous tissues found in one database but not the other ^[8-10]^. The proportion of each tissue assignment may depend on the number of markers used for each. Because of this, the differences between the cfDNAme and Meth-Atlas libraries on the order of thousands for some tissues could lead to estimated tissue proportion differences seen between the two methods. In fact, cfDNAme tends to estimate higher proportions of erythrocyte progenitor cells in each gold-standard sample, as it has fewer markers (several of which overlap with Neutrophil markers) to correctly determine tissue type. One fix for this would be to use a subsampled cfDNAme database, using the top 300 (or fewer) marker windows, which may improve the database similarities. Additionally, adding more samples for erythrocyte progenitor cells to improve the markers used may also help.

Overall, here, we believe we have demonstrated that, given adequate coverage of a sample, a methylome array-based deconvolution method is as successful for tissue of origin identification in cfDNA as a WGBS based method, and in some cases may be appropriate to use either by themselves, or in combination with a WGBS based method to improve the confidence of tissue of origin assignments in cfDNA samples.

