## Supplementary Figures for "Benchmarking and optimization of cell-free DNA deconvolution"

**Supplementary Figure 1: Validation of cfDNAme deconvolution algorithm.** (A) Genomic DNA (gDNA) mixtures from neutrophil and human vascular endothelial cells. (B) Correlation between DNA methylation and automated hematological analyzer for leukocyte differential (granulocytes, lymphocytes, and monocytes) estimate.

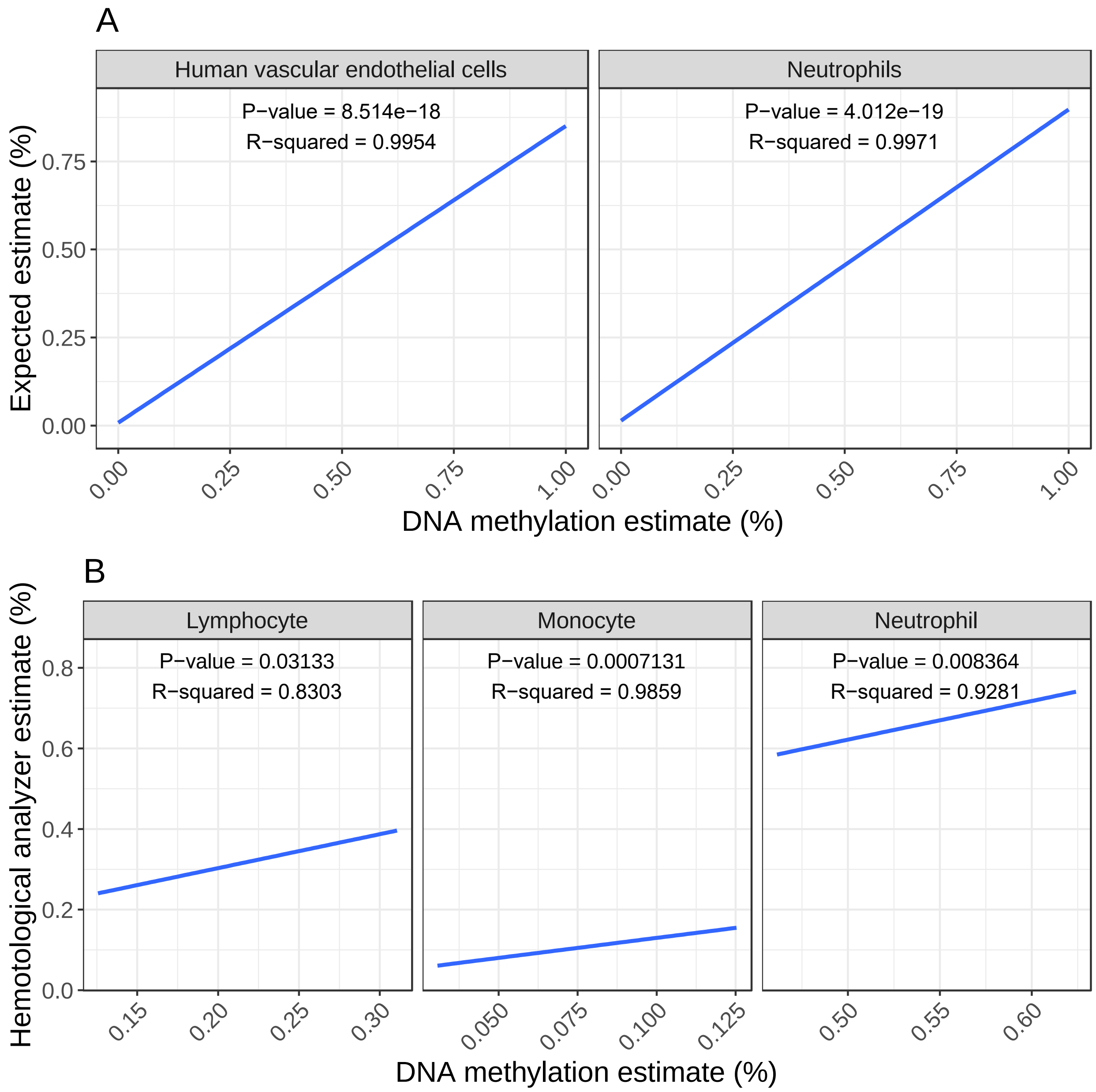

**Supplementary Figure 2:** Association between CpG specificity/effect size and the prevalence/frequency in a group. The blue line shows the (significant) correlation between the two values in each group.

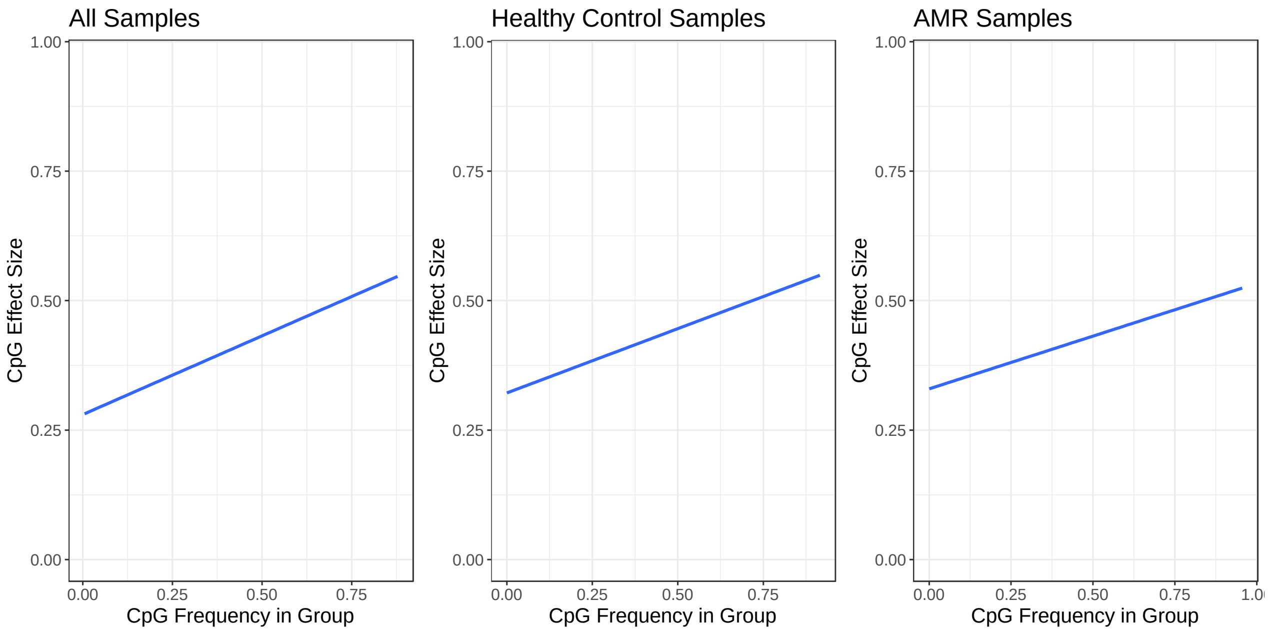

**Supplementary Figure 3:** Overlap of CpG markers found in healthy control and AMR groups, for the top 100, 500, 1000, 2000, 5000 & 7000 CpG markers where markers are ranked by their specificity of methylation state to a tissue. Note that though we looked at the top CpG markers based on specificity to a tissue, there is no guarantee these CpG markers are present in our assessed samples, resulting in only ~60% of CpGs being shown in the Venn diagrams.

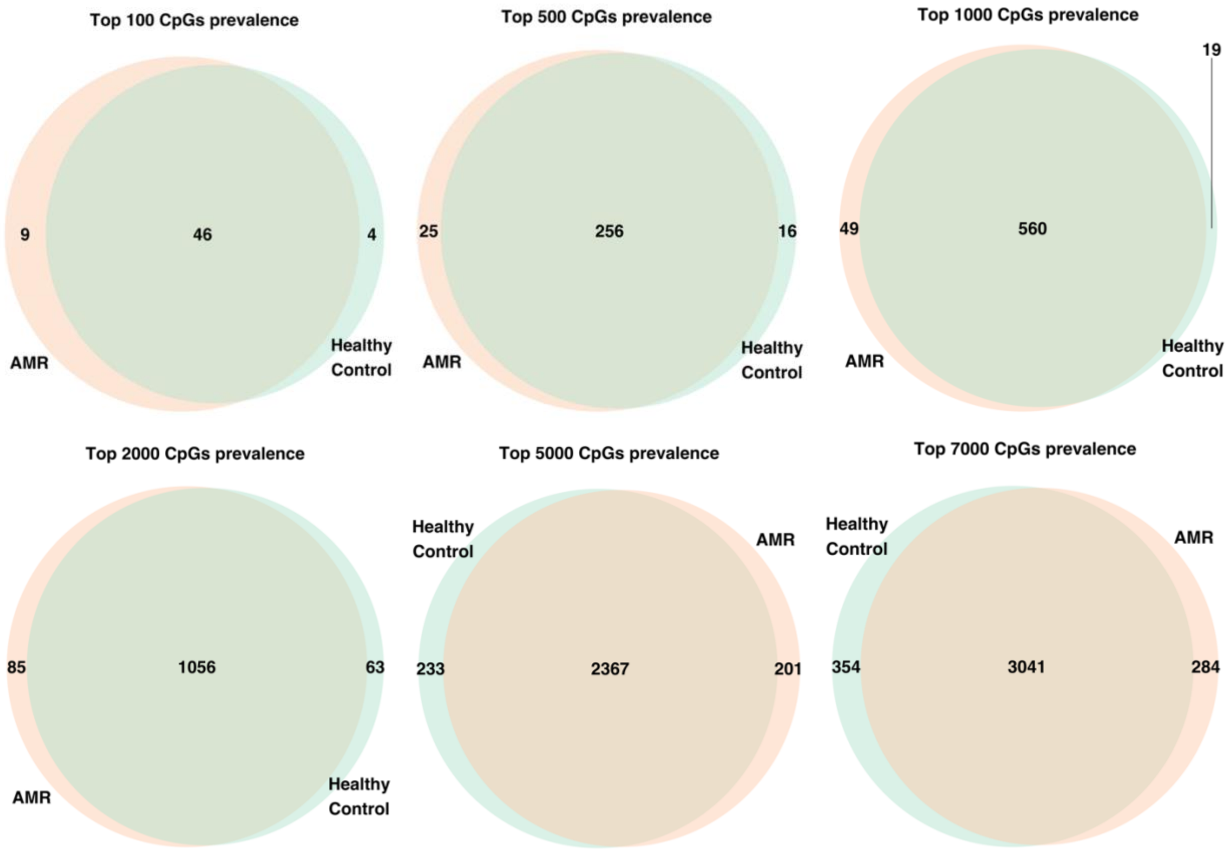

**Supplementary Figure 4:** Comparison of cfDNAme & Meth-Atlas deconvolution results for each sample at full coverage.

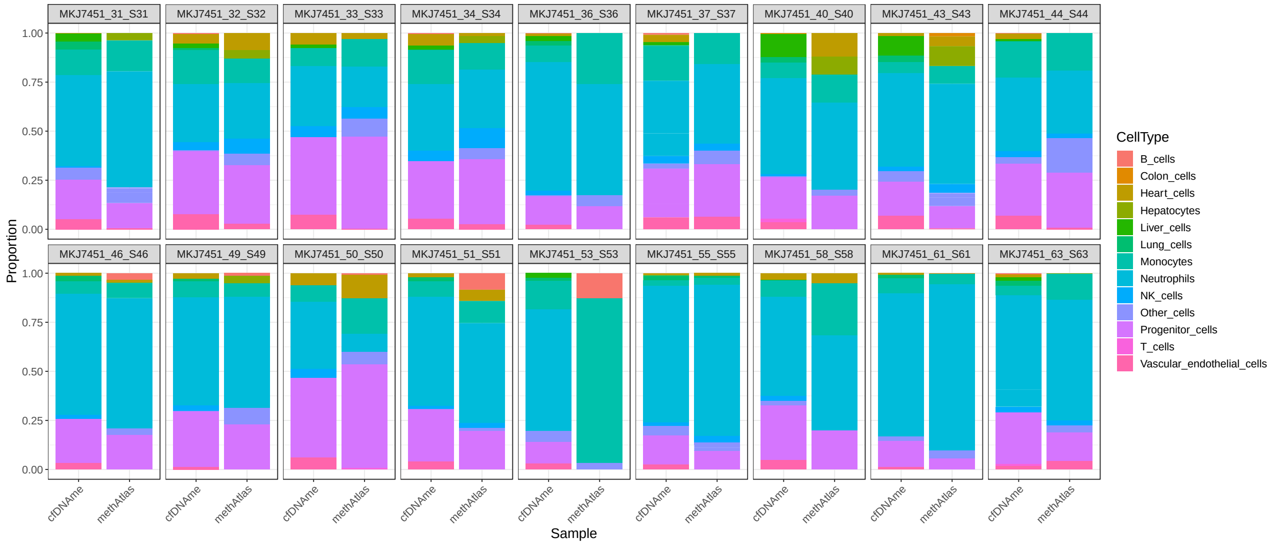

**Supplementary Figure 5:** Overlapping number of CpGs found between Meth-Atlas cell type categories, shown on a logarithmic scale, note the high amount of overlapping CpGs between Neutrophils, Monocytes and Progenitor cells.

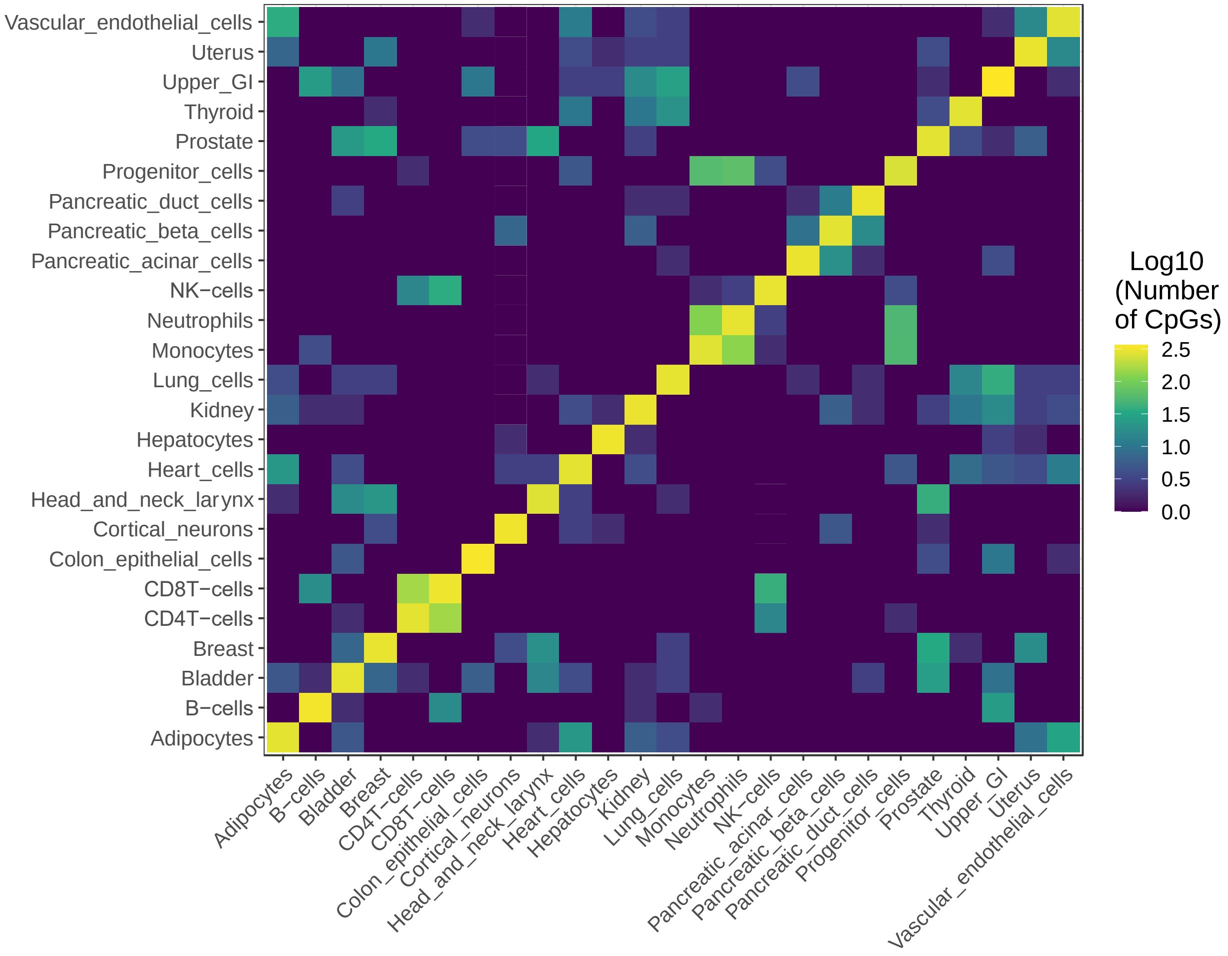

**Supplementary Figure 6:** Deconvolution tissue type proportions for each sample, separated by sequencing lane. Proportions are estimated from cfDNAme and Meth-Atlas. Note that due to their small proportion, we have merged Uterus cells, Gastrointenstinal cells, Bladder cells, Pancreas cells, Breast cells, Neuron cells, Adipose cells, and Skin cells as ‘Other cells’.

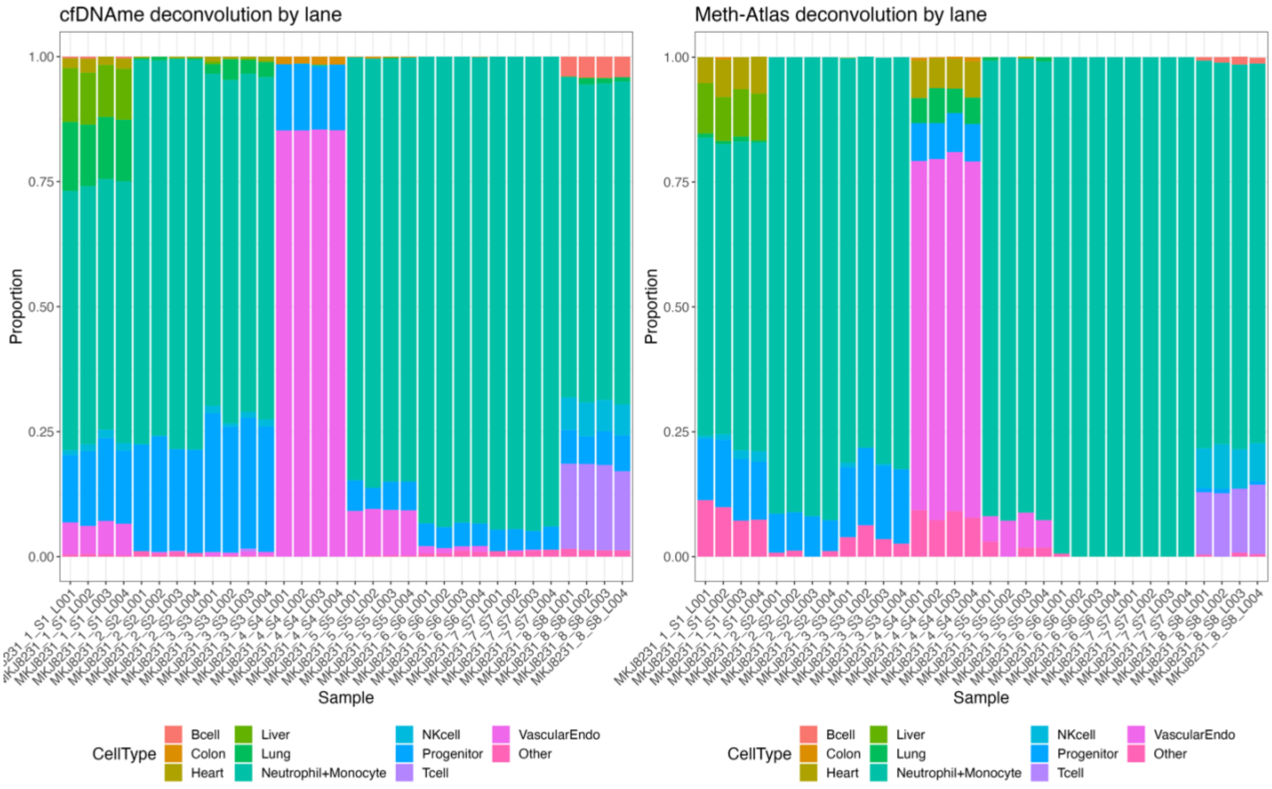

**Supplementary Figure 7:** Median cell type proportions across replicates at different subsampled coverage proportions, for each sample, estimated using cfDNAme and Meth-Atlas. Stacked barplots are colored by the proportion of each tissue. Note that due to their small proportion, we have merged Uterus cells, Gastrointenstinal cells, Bladder cells, Breast cells, Neuron cells, Adipose cells, and Skin cells as 'Other cells'.

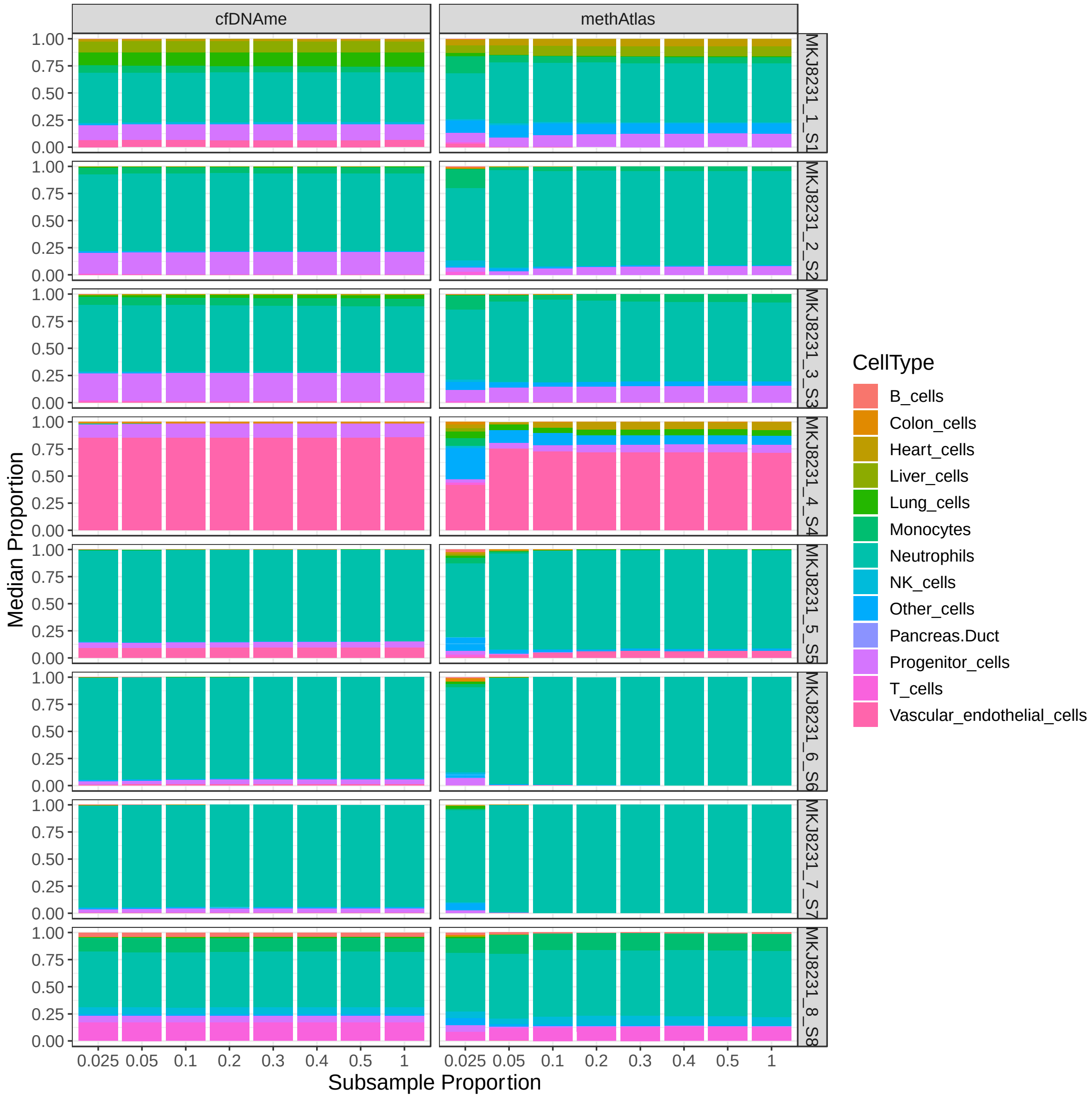

**Supplementary Figure 8:** Number of CpGs seen in our gold standard subsampled datasets, with samples grouped by the proportion of reads used in each subsampled dataset.

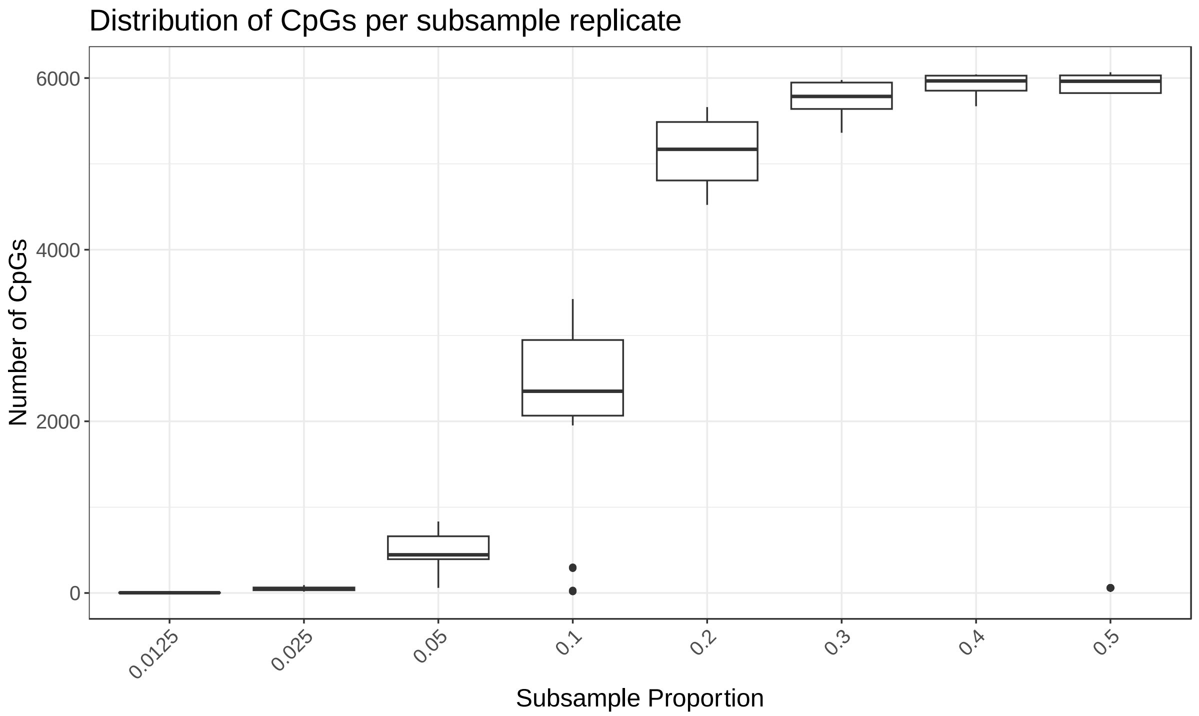
